# Evolving mutation rate advances invasion speed of sexual species

**DOI:** 10.1101/008979

**Authors:** Marleen M. P. Cobben, Alexander Kubisch

## Abstract

Many species are shifting their ranges in response to global climate change. The evolution of dispersal during range expansion increases invasion speed, provided that a species can adapt sufficiently fast to novel local conditions. Mutation rates can evolve too, under conditions that favor an increased rate of adaptation. However, evolution at the mutator gene has thus far been deemed of minor importance in sexual populations due to its dependence on genetic hitchhiking with a beneficial mutation at a gene under selection, and thus its sensitivity to recombination. Here we use an individual-based model to show that the mutator gene and the gene under selection can be effectively linked at the population level during invasion. This causes the evolutionary increase of mutation rates in sexual populations, even if they are not linked at the individual level. The observed evolution of mutation rate is adaptive and clearly advances range expansion both through its effect on the evolution of dispersal rate, and the evolution of local adaptation. In addition, we observe the evolution of mutation rates in a spatially stable population under strong directional selection, but not when we add variance to the mean selection pressure. By this we extend the existing theory on the evolution of mutation rates, which is generally thought to be limited to asexual populations, with possibly far-reaching consequences concerning invasiveness and the rate at which species can adapt to novel environmental conditions as experienced under global climate change.

## Introduction

Many species are currently expanding their ranges as a response to increasing global temperatures under climate change (Chen et al. 2011). Range expansions are known to have profound effects on the genetic composition of populations, regarding both neutral and adaptive genetic diversity (Hewitt 1996; Phillips et al. 2006; Excoffier et al. 2009; Cobben et al. 2012b). Traits that act to increase species’ dispersal capabilities and population growth rates are selected for under range expansions due to spatial sorting (Hill et al. 2011; Shine et al. 2011) and kin competition (Kubisch et al. 2013b). This may lead to higher dispersal rates (Thomas et al. 2001; Kubisch et al. 2010), dispersal distances (Phillips et al. 2006) and effective fertilities (Moreau et al. 2011) at the expanding front of species’ ranges due to micro-evolution. An increasing dispersal rate under range expansion will increase the invasion speed (Phillips et al. 2006), but only if the species is able to adapt sufficiently rapid to novel local conditions. However, under range expansion the depletion of genetic diversity at the expanding range border due to iterated founder effects (Hewitt 1996; Excoffier et al. 2009; Cobben et al. 2011) can be expected to limit the invasion speed as low genetic diversity will lead to low rates of local adaptation and thereby delayed population establishment.

Evolvability, i.e. a set of mechanisms that facilitates evolution, has been shown to be subjected to selection under conditions that favor an increased rate of local adaptation, e.g. under increasing environmental stochasticity and stress (Earl and Deem 2004; Kashtan et al. 2007; Lee and Gelembiuk 2008). One example is the evolution of mutation rates, where selection can act on allelic variation in the processes of DNA repair and as such result in increased mutation rates, which cause higher levels of genetic diversity and thus enable adaptation to changing selection pressures (Kimura 1967; Leigh Jr 1970; Leigh Jr 1973; Sniegowski et al. 1997; Taddei et al. 1997; Metzgar and Wills 2000; Sniegowski et al. 2000; Bedau and Packard 2003). During the colonization of a spatially heterogeneous environment, an increased mutation rate will be selected for as it generates more genetic diversity, provided that this results in the faster establishment of new populations by pre-adapted individuals. Further increased selection for high mutation rates can be expected due to spatial sorting (Shine et al. 2011) and increased relatedness between individuals (Kubisch et al. 2013b) at the expansion front, in combination with the increased dispersal rates and higher potential invasion speed (Phillips et al. 2006; Phillips et al. 2010a). With the establishment of a stable range border, after range expansion, the selection pressures change and a return to lower mutation rates is expected.

The evolution of mutation rates has thus far mostly been associated with and only shown for asexual populations (Drake et al. 1998; Sniegowski et al. 2000; Raynes et al. 2011). As selection acts on the mutation that occurs at a gene under selection and not on the rate with which such mutations occur, the evolution of mutation rates is the result of indirect selection. The establishment of a particular mutation rate is thus restricted to genetic hitchhiking, which is highly sensitive to recombination (Drake et al. 1998; Sniegowski et al. 2000). However, here we hypothesize that these two genes (the mutator gene and the gene under selection) need not be linked at the individual level for an increase of the mutation rate to occur, if they are sufficiently linked at the population level. If an individual can only mate with genetically similar individuals, genetic information at the mutator gene, i.e. the DNA repair gene, and the gene under selection can be reunited in the offspring despite their independent inheritance, in a form of genetic drift. Such conditions of genetic similarity are typical for range expansions due to founder effects at the expanding range margin, or in systems under strong directional selection. Here we investigate whether the evolution of mutation rates can contribute to the adaptive potential of sexual populations under such conditions.

In this study, we use a spatially explicit individual-based metapopulation model of a sexual species establishing its range on a spatial gradient to investigate 1) whether mutation rates increase during range expansion, 2) if this is related to the rate of dispersal during range expansion, and 3) if this is related to the directionality of the selection. The results show the co-evolution of dispersal rates and mutation rates in sexual populations resulting from the particular properties of spatial sorting. This co-evolution leads to a faster range establishment, and has possibly far-reaching consequences concerning our knowledge of invasiveness and the rate at which species can adapt to novel environmental conditions.

## Methods

We use a spatially explicit individual-based metapopulation model of a sexually reproducing species with discrete generations, inspired by insects’ ecology and parameterized using empirical data (Poethke et al. 1996; Amler et al. 1999), which expands its range along a gradient in temperature. We allow the mutation rate to evolve, and investigate its interplay with the evolution of dispersal rate and temperature adaptation during and after range establishment.

### Landscape

The simulated landscape consists of 250 columns (*x*-dimension) of 20 patches each (*y*-dimension). We assume wrapped borders in y-direction, building a tube. Hence, if an individual leaves the world in y-direction during dispersal, it will reenter the simulated world on the opposite side. However, if it leaves the world in the *x*-direction, it is lost from the simulation. To answer our research questions the model requires a need for local adaptation during range expansion. Thus every column of patches (*x*-position) is characterized by its specific mean temperature *τ_x_*. This mean local temperature is used for the determination of the level of local adaptation of individuals. To simulate a large-scale habitat gradient, *τ_x_* changes linearly from *τ_x_*_=1_ = 0 to *τ_x_*_=250_ = 10 along the x-dimension, i.e. by Δ*_τ, x_* = 0.04 when moving one step in *x*-direction.

### Population dynamics and survival of offspring

Local populations are composed of individuals, each of which is characterized by several traits: 1) its sex, 2) its dispersal probability determined by the alleles at the dispersal locus *l_d_*, 3) its optimal temperature *τ_opt_*, i.e. the temperature under which it survives best, determined by the alleles at its adaptation locus *l_a_* (see below for details), 4) its genetic mutation rate determined by the alleles at the mutator locus *l_m_* (see below under Genetics), and 5) a diploid neutral locus *l_n_*, for the sake of comparing the levels of genetic diversity with other loci.

Local population dynamics follow the time-discrete Beverton–Holt model (Beverton and Holt 1957). Each individual female in patch x, y is therefore assigned a random male from the same habitat patch (males can potentially mate several times) and gives birth to a number of offspring drawn from a Poisson distribution with mean population growth rate *λ*. The offspring’s sex is chosen at random. Density-dependent survival probability *s*_1_ of offspring due to competition is calculated as:

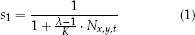

with *K* the carrying capacity and *N*_*x, y, t*_ the number of individuals in patch *x, y* at time *t*. Finally, the surviving offspring experience a further density-independent mortality risk (1 – *s*_2_) that depends on their local adaptation, so the matching of their genetically determined optimal temperature (*τ_opt_*) to the temperature conditions in patch *x, y* (*τ_x_*) according to the following equation:

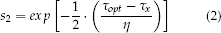

where *η* describes the niche width or ‘tolerance’ of the species. We performed simulations for the species with a niche width of *η* = 0.5, equivalent to a decrease of survival probability of about 0.02 when dispersing one patch away from the optimal habitat. In this approach we assume that density-dependent mortality (1 – *s*_1_) acts before mortality due to maladaptation to local conditions (1 – *s*_2_). In addition, each population has an extinction probability *∊* per generation. Individual surviving offspring disperse with probability d that is determined by their dispersal locus (see below). If an individual disperses it dies with probability *μ*, which is0.2 throughout the landscape. This mortality accounts for various costs that may be associated with dispersal in real populations, like fertility reduction or predation risk (Bonte et al. 2012). We assume nearest-neighbor dispersal, i.e. successful disperses settle randomly in one of the eight surrounding habitat patches.

### Genetics

As mentioned above, each individual carries three unlinked, diploid loci coding for its dispersal probability, its optimum temperature (and thus its degree of local adaptation), and its genetic mutation rate, respectively, and an additional neutral locus. The phenotype of an individual is determined by calculating the means of the two corresponding alleles, with no dominance effect involved. Hence, dispersal probability d is given by 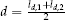 (with *l_d_*,_1_ and *l_d_*,_2_ giving the two ‘values’ of the two dispersal alleles), optimal temperature *τ_opt_* is calculated as *τ_opt_* 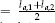 (with *l_a_*,_1_ and *l_a_*,_2_ giving the ‘values’ of the two adaptation alleles), and similarly the mutation rate *m* = 10^−exp^ (with *exp* 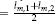, and *l_m_*,_1_ and *l_m_*,_2_ the ‘values’ of the two mutator alleles). At each of the four loci, newborn individuals inherit alleles, randomly chosen, from the corresponding loci of each of their parents. During transition from one generation to the next an allele may mutate with the genetically determined probability *m* given by the value based on the two alleles at the mutator locus *l_m_* as elaborated above. Mutations are simulated by adding a random number to the value of the inherited allele. This value is drawn from a Gaussian distribution with mean 0 and standard deviation 0.2. The lethal mutation probability is Ω = 0.1 however, so tern percent of the mutations cause immediate death of the individual.

### Simulation experiments

Simulations were initialized with a ‘native area’ (from *x* = 1 to *x* = 50) from where the species was able to colonize the world, while the rest of the world was initially kept free of individuals. Upon initialization, dispersal alleles (*l_d, i_*) were randomly drawn from the interval 0 < *l_d, i_* < 1, and mutator alleles *l_m, i_* were set to 4, which set the initial mutation rate *m* to 10^−4^. Populations were initialized with *K* locally optimally adapted individuals, i.e. adaptation alleles were initialized according to the local temperature *τ_x_*. However, to account for some standing genetic variation we also added to every respective optimal temperature allele a Gaussian random number with mean zero and standard deviation 0.2. Identical copies of these alleles were used to initialize the neutral locus as well, for sake of comparison. We performed 200 replicate simulations, which all covered a time span of 15,000 generations. To establish equilibrium values, the individuals were confined to their native area during the first 10,000 generations. After this burn-in period, the species was allowed to pass the *x* = 50 border and expand its range for the remaining 5,000 generations. Table 1 summarizes all relevant model parameters, their meanings and the standard values used for the simulations.

**Table 1.** Parameter values.

| parameter / variable | (initialization) value | meaning |
| --- | --- | --- |
| *individual parameters:* |  |  |
| $l_{d,1}, l_{d,2}$ | (0 to 1 and) evolving | alleles coding for dispersal propensity |
| $l_{a,1}, l_{a,2}$ | (optimal with sd 0.5 and) evolving | alleles coding for optimal temperature |
| $l_{m,1}, l_{m,2}$ | (4 and) evolving | alleles coding for mutation rate of optimal temperature |
| $l_{n,1}, l_{n,2}$ | (0.5 with sd 0.5 and) evolving | neutral alleles as control |
| *simulation parameters:* |  |  |
| $K$ | 100 | carrying capacity |
| $\lambda$ | 2 | per capita growth rate |
| $\epsilon$ | 0.05 | local extinction probability |
| $\Omega$ | 0.1 | lethal mutation probability |
| $m$ | $10^{-4}$ | mutation rate for dispersal and evolvability alleles |
| $\mu$ | 0.2 | local dispersal mortality |
| $\tau_x$ | [0..10] | local temperature |
| $\eta$ | 0.5 | niche width |
| $x_{max}$ | 250 | extent of simulated landscape in $x$-direction |
| $y_{max}$ | 50 | extent of simulated landscape in $y$-direction |

Several control simulations were performed:

1. To determine whether the evolution of mutation rates was neutral or adaptive. For this, the simulations were repeated with 200 replicates for 1. fixed values of dispersal rate, *d* = 0.05, *d* =0.1 and *d* =0.2, while allowing the mutation rate to evolve, 2. with fixed values of the mutation rate *m* of 10^−4^ and 10^−5^, combined with evolving dispersal rate, and 3. with both rates fixed, investigating the combination of *d* =0.2 and *m* = 10^−4^ and of *d* =0.2 and *m* = 10^−5^.
2. A control simulation was performed with 90 percent lethal mutations.
3. To assess the role of repeated colonizations and directional selection for the evolution of mutation rates we performed control simulations under a variable spatial gradient in temperature. For this we applied a new habitat gradient, where *τ_x_* still changes from *τ*_*x* = 1_ = 0 to *τ*_*x* =250_ = 10 along the *x*-dimension, but at each *τ_x_* we added a random number in the range [–0.5,0.5]. In addition we simulated a non-expanding population under both temperature gradients, so applying a non-variable and variable temporal gradient in temperature. For these last experiments the species was initialized in the whole landscape, so not restricted to the native area, and the temperature was steadily increased in the entire landscape. With this equal selection pressure for the species was forced, but without a range expansion, by changing the temperature at the same rate as experienced by the marginal populations in the spatial scenarios. Global dispersal was applied here.

### Analysis

The individual phenotypes for the three traits were documented in time and space throughout the simulations and averaged per *x*-position. For the dispersal rate we calculated the arithmetic mean, while the mutation rate was averaged geometrically because the mutator gene codes for the exponent’s value. Genetic diversity was calculated as the variance in allelic values at the adaptation locus, the dispersal locus and the neutral locus, per *x*-position.

## Results

After the burn-in phase, the species’ range expands across the landscape (Fig. 1). The landscape is fully colonized after between 1,000 and 1,500 generations (Fig. 1A). During range expansion the average dispersal rates and mutation rates increase, and they decrease again after the colonization is complete and the range border has stabilized (Fig. 1B/C). Genetic diversity at the different loci shows a typical pattern of range expansion (due to founder effects or spatial sorting) with little genetic diversity at the expanding range margin, which increases with the age of the populations (Fig 1D-F). The maximum level of genetic diversity differs between the different loci.

**Figure 1.**
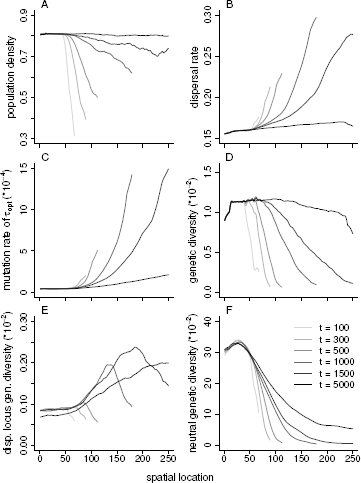
The average values over 200 simulations during and after range expansion across the gradient (horizontal axis) in time (gray scaling from light to dark, as time proceeds, which is given in a sequence of generations 100, 300, 500, 1000, 1500, 5000) of A. population density, B. dispersal rate, C. the mutation rate, D. genetic diversity at the adaptation locus, E. genetic diversity at the dispersal locus, and F. neutral genetic diversity, all measured as the variance in allele values. For reasons of clarity, a moving average with a window size of 20 has been applied (data were present in 10-generation intervals).

The control simulations with fixed rates (Fig. 2) show that a fixed, lower mutation rate causes a lower speed of invasion, both for evolving dispersal rates (Fig. 2a panel A) as for fixed dispersal rate (Fig. 2a panel B). Under fixed dispersal rates (Fig. 2b) we see the evolution of higher mutation rates and faster range expansions with higher dispersal rates. The control simulations in which 90 percent of the mutations were lethal (Ω = 0.9) also show an increase of the dispersal and mutation rates (Fig. S1) but to a lesser extent than in the original simulations.

**Figure 2.**
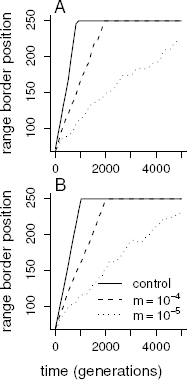
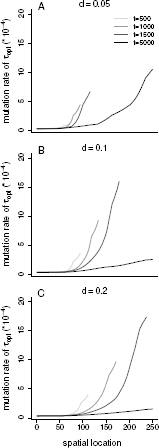
The results of the control simulations, where in a. the range border position in time (horizontal axis) is shown, averaged over 200 simulations for the original experiment (with evolving dispersal rate) in panel A for the case with evolving mutation rate (‘control’) and fixed mutation rates of 10^−4^ and 10^−5^. Panel B is the same, but for a fixed dispersal rate of 0.2. In b. the average values of the mutation rate during and after range expansion across the gradient (horizontal axis) is shown in time (gray scaling from light to dark, as time proceeds, which is given in a sequence of generations 500, 1000, 1500, 5000) under A. a fixed dispersal rate of 0.05, B. a fixed dispersal rate of 0.1, and C. a fixed dispersal rate of 0.2.

For the spatially stable population a strongly directional selection showed an increase of dispersal and mutation rates in time (Fig. 3a, B/C), while the mutation rate could not evolve under variable temperature increase (Fig. 3b, C). This is in contrast to the expanding population, in which the mutation rate could increase while expanding across the variable temperature gradient in space (Fig. 4C). The local level of adaptation *s*_2_ is close to one in all simulations, throughout the simulation time and across the entire species’ range.

**Figure 3.**
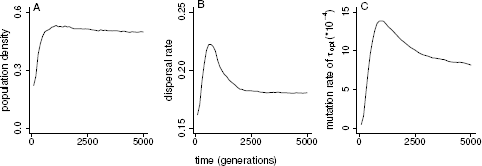
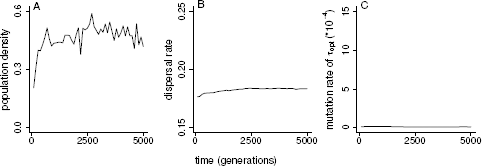
The average values over 200 simulations in time in a population subjected to a. a temporal non-variable gradient in temperature, and b. temporal variable gradient in temperature, of A. population density, B. dispersal rate, and C. the mutation rate. For reasons of clarity, a moving average with a window size of 20 has been applied (data were present in 10-generation intervals).

**Figure 4.**
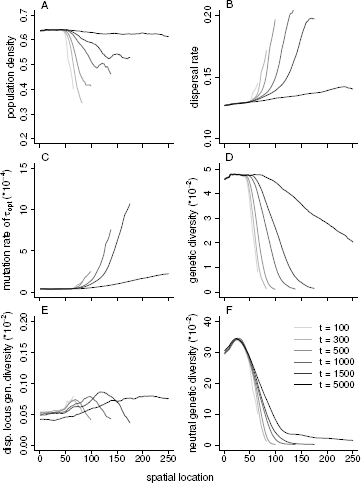
Applying a variable spatial gradient in temperature, the average values are given over 200 simulations during and after range expansion across the gradient (horizontal axis) in time (gray scaling from light to dark, as time proceeds, which is given in a sequence of generations 100, 300, 500, 1000, 1500, 5000) of A. population density, B. dispersal rate, C. the mutation rate, D. genetic diversity at the adaptation locus, E. genetic diversity at the dispersal locus, and F. neutral genetic diversity, all measured as the variance in allele values. For reasons of clarity, a moving average with a window size of 20 has been applied (data were present in 10-generation intervals).

## Discussion

In this study we investigate whether mutation rates can evolve upwards under the range expansion of a sexual species that needs to adapt to novel local temperature conditions. We observe an increase of the mutation rate, which leads to a faster evolution of dispersal rates and thus faster range expansion. This also occurs when we apply variance to the mean temperature gradient in space, and, to a lesser extent, even when we assume that 90 percent of the mutations are lethal. Under fixed dispersal rates we also observe the evolution of mutation rates with a similar advancing effect on the invasion speed due to the faster rate of local adaptation. The increase of the mutation rate occurs despite the independent inheritance of the three traits in sexual populations due to the particular properties of spatial sorting during range expansion. In a spatially stable population subjected to a temporal increase of temperature the mutation rate evolves as well, but not when variance is added to the mean temperature increase.

During range expansion the dispersal rate shows a clear signal of spatial sorting (Shine et al. 2011) and kin competition (Kubisch et al. 2013b), with good dispersers gathering at the expanding wave front (Phillips et al. 2010a). A high mutation rate is beneficial since it allows the faster occurrence of alleles coding for higher dispersal rates that establish on the expansion wave and increase the invasion speed. From the results with fixed dispersal rates it shows that a high mutation rate is also beneficial as it causes the faster occurrence of alleles at the gene for local adaptation to temperature. In both cases the individuals carrying alleles for high mutation rates are the first to establish new populations because they carry these beneficial and novel alleles at the dispersal and adaptation loci as well. This result is particularly interesting as selection for optimum mutation rate is associated with asexual populations (Kimura 1967; Leigh Jr 1970; Leigh Jr 1973; Sniegowski et al. 2000). Indeed, selection only operates on the dispersal and adaptation loci, favoring mutations that increase the speed of range expansion. In sexual populations, strong linkage is required for the (advantageous) alleles at the these loci and the (high) mutation rate allele at the mutator locus to be inherited together, and as such to lead to indirect selection at the mutator locus (Johnson 1999; Sniegowski et al. 2000; Tenaillon et al. 2000). In our study, however, these two loci are genetically unlinked. Yet, the colonization of an empty patch on average occurs by individuals carrying high mutation rate alleles. A high initial population growth rate and the non-random set of invaders in such a newly established population result in a local non-random subset of the available genetic variation at the mutator locus. As such the beneficial alleles at the adaptation and dispersal loci, and high mutation rate alleles at the mutator locus are here essentially ‘soft-linked’ for lack of availability of low mutation rate alleles. This effective link at the population level can apparently substitute a genetic link at the individual level. In the spatially stable population under variable selection pressure we see that this link is broken in what could be called the population-level equivalent of recombination.

The different functions of the genes are reflected in the effects of high mutation rates. At the neutral locus genetic diversity steadily increases, towards a maximum determined by population dynamics. At the gene that determines the individual’s optimal temperature the maximum genetic diversity is determined by the number of allele values that allow the individual’s survival at that particular local temperature. Comparing Fig.1D and Fig. 4D shows the difference between the variable and non-variable temperature gradient. Mutation rates above a certain threshold do not add more genetic diversity at these two loci. This is in contrast to the dispersal locus, where a higher mutation rate causes a higher level of genetic diversity. Both high rate signals of dispersal and mutation disappear with time, as anticipated. High dispersal rates are only favorable with frequent population extinctions and low dispersal mortality (Ronce 2007). Once the range border stabilizes a low dispersal phenotype is more advantageous due to the assumed dispersal mortality. However, these slow dispersers by definition take some time to reach the area (genetic signature of range expansion, Phillips et al. 2010b; Cobben et al. 2015), especially when the mutation rate levels off and new dispersal phenotypes only slowly appear locally. The decrease of the mutation rate is also caused by the processes at the temperature locus. Once the maximum level of genetic diversity has been reached here, and population densities are at their equilibrium values, more mutations cause maladaptation and lower levels of the mutation rate are more beneficial.

In relation to this last point, we have modeled the mutation rate as the probability that an inherited allele mutates. Since these mutation rates are caused by genes that are involved in processes of DNA repair (Metzgar and Wills 2000), high mutation rates will however likely affect the individual itself and not only its offspring. This is the result of mutations occurring when DNA is copied during the division of cells other than only the reproduction cells. Such mutations might then cause defects or tumors. Modeling mutation rates that negatively affect the individual’s fitness will likely change our results, because high mutation rates are then more disadvantageous (Kimura 1967; Leigh Jr 1970). On the other hand, there are large differences in mutation rates between (parts of) genomes (Drake et al. 1998) and DNA repair is not restricted to a single pathway (Rottenberg et al. 2008). In addition, if a high mutation rate only affects an individual’s survival after reproduction, then high mutation rates might evolve despite their negative effects for the individual.

Our results can be affected by the used genetic architecture, where linkage between traits (Blows and Hoffmann 2005; Hellmann and Pineda-Krch 2007), polygeny, and the magnitude of mutations can be of importance in range dynamics (Kawecki 2000, 2008; Gomulkiewicz et al. 2010). The used mutation model of adding values to the inherited values result in mutations that are at most mildly deleterious at the adaptation locus, while the distribution of random mutations would invoke a stronger selection pressure (Sanjuán et al. 2004). In addition, the use of an infinite allele model for mutations would allow the occurrence of extreme dispersal values at any mutation event, which would probably reduce the maximum mutation rate values. However, our results are based on the assumption that ten percent of the mutations are lethal and we still see a significant increase of the mutation rate when we assume 90 percent lethal mutations (Fig. S1). This is in contrast to what was found in an experimental study of sexual populations of yeast (Raynes et al. 2011), which has however not taken a spatial perspective.

We observe that mutation rate can increase in combination with the increased dispersal rates and spatial variation, as experienced under range expansion. High dispersal rates, i.e. the immigration of many individuals is expected to maintain a high local level of genetic variation (Holt and Barfield 2011), from which one would expect high levels of dispersal to be accompanied by a low local mutation rate. At the margin, however, relatedness amongst individuals increases at an advancing range front (Kubisch et al. 2013b), reducing both local genetic diversity and the diversity of immigrants. Under these conditions an increase in the mutation rate evolves, which allows faster adaptation to the experienced spatial variation in local temperature, causing a faster range expansion across the spatial gradient.

Holt and Barfield (2011) investigated niche evolution at species’ range margins and found that local evolution is hampered when source populations of immigrating individuals are at low density, as a result of the stochastic processes in such populations (Pearson et al. 2009; Bridle et al. 2010; Turner and Wong 2010). The likelihood of observing niche evolution is further affected by the mutation rate, where dispersal limits local evolution in the sink population under a higher mutation rate, because of the increased numbers of maladapted individuals from the source (Holt and Barfield 2011). They did, however, not allow the joint evolution of mutation rate and dispersal rate, but instead used fixed rates. As a result, the dispersal rate does not decrease after colonization, while the conditions in the sink population make its persistence dependent on the constant influx of (maladapted) individuals, both in contrast to the model presented here.

In our study we investigate the evolution of mutation rates. Dealing with novel environmental conditions or increased evolvability is however not restricted to mutation rates, but can be modeled in different ways, e.g. an increased magnitude of the phenotypic effect of mutations (Griswold 2006), an epigenetic effect, the evolution of modularity (Kashtan et al. 2009), degeneracy (Whitacre and Bender 2010), or the evolution of generalism or plasticity (Lee and Gelembiuk 2008). In addition, Kubisch et al. (2013b) showed that when dispersal is a means of adaptation, by tracking suitable conditions during periods of change, genetic adaptation does not occur. Which kind of adaptation to change can be expected under specific ecological and environmental conditions is an interesting field of future investigation.

There is an ever-expanding pool of literature discussing the ecological and evolutionary dynamics of dispersal in the formation of species ranges (reviewed in Kubisch et al. 2014). While individual-based models have recently largely extended our theoretical knowledge of interactions and evolution of traits during range expansion, empirical data have been restricted to a few well-known cases (Thomas et al. 2001; Phillips et al. 2006; Moreau et al. 2011; Fronhofer and Altermatt 2015). Increasing ecological realism in our models (Cobben et al. 2012a) can improve the predictability of theoretical phenomena which can then be tested by data from field studies. So far, increased dispersal has been shown to increase invasion speeds (Thomas et al. 2001; Phillips et al. 2010a), affect the fate of neutral mutations (Travis et al. 2010), as well as the level of local adaptation (Kubisch et al. 2013a), and local population dynamics (Ronce 2007), and in addition causes strong patterns of spatial disequilibrium (Phillips et al. 2010b; Cobben et al. 2015).

In this study we show new and unexpected consequences of the particular genetic properties of populations under spatial disequilibrium, i.e. the co-evolution of dispersal rates and mutation rates, even in a sexual species and under realistic spatial gradients, resulting in faster invasions. We conclude that range expansions and the evolution of mutation rates are in a positive feedback loop, with possibly far-reaching ecological consequences concerning invasiveness and the adaptability of species to novel environmental conditions.

## Acknowledgements

We thank Troy Day, Andy Gardner, Marcel Visser, Thomas Hovestadt, and René Smulders for their encouragement and valuable comments that have contributed to the improvement of this manuscript. This work was supported by the Open Program of the Netherlands Organization of Scientific Research (NWO 822.01.020 for M.M.P.C.), and the German Science Foundation (KU 3384/1-1 for A.K.).

